# Differential Contributions of Anterior Cingulate and Orbito-Frontal Cortex to action timing and its self-monitoring in rats

**DOI:** 10.64898/2026.03.25.714178

**Authors:** Lea Le Barillier, Valerie Doyeere, Tadeusz W. Kononowicz

## Abstract

Adaptive behavior requires the ability to monitor the accuracy of self-generated actions, including the production of precise time intervals. Here, we investigated the neural mechanisms underlying temporal error monitoring in rats by combining a time production and error-reporting task with selective pharmacological inactivation of the orbitofrontal cortex (OFC) and anterior cingulate cortex (ACC). We found that OFC inhibition impaired the production of time intervals in a dose-dependent manner, indicating its critical role in generating temporally precise actions, whereas ACC inhibition left time production intact but caused a systematic overestimation of temporal errors and increased overconfident responses on incorrect trials. Analyses of choice behavior revealed that ACC inactivation disrupted the use of trial history and shifted decision thresholds, suggesting that ACC implements a hierarchical read-out of ongoing temporal performance. These results support a functional dissociation in which OFC provides the temporal signal for action, while ACC evaluates errors and confidence. Our findings establish a causal link between prefrontal circuits and self-monitoring of time, offering a model for hierarchical temporal control and evaluation in the rodent brain.

## Introduction

The ability to evaluate the accuracy of one’s own behavior is a fundamental aspect of adaptive cognition. Such self-monitoring allows organisms to assess internal computations and adjust behavior in the absence of external feedback (Dehaene et al., 2017). In parallel, the ability to generate and control behavior in time is a core function of living systems. Together, these capacities raise the question of how the brain monitors errors arising from self-generated timing behavior, such as the production of precise time intervals.

Previous work from our group (Kononowicz et al., 2022) established that rats are capable of monitoring their own timing errors. In that study, animals produced time intervals and subsequently categorized their performance as small or large errors, reliably reporting this distinction when given a choice. These findings demonstrate that rats can form an internal estimate of their temporal accuracy, providing a robust behavioral framework to investigate the underlying neural mechanisms.

Among candidate substrates, two prefrontal regions stand out: the orbitofrontal cortex (OFC) and the anterior cingulate cortex (ACC). Both areas have been broadly implicated in performance monitoring, decision-making, and evaluation under uncertainty (Cohen et al. 2005; Wallis et al. 2011), yet their specific contributions remain debated. The ACC has been consistently associated with error detection and action monitoring, with studies showing that it encodes motor errors (Totah et al. 2019; Brockett et al. 2020), contributes to confidence judgments (Stolyarova et al. 2019), and processes feedback to guide behavioral adaptation (Wallis et al. 2011; Hayden et al. 2011). In parallel, the OFC has been linked to value-based evaluation and confidence estimation, with impairments in similar tasks following its disruption (Kepecs et al. 2008; Lak et al. 2014; Masset et al. 2020). Importantly, the OFC has also been implicated in action timing and the representation of temporal structure (Cazares et al. 2022), suggesting a potential role at the interface between timing and evaluation. Despite this overlap, it remains unclear whether these regions contribute similarly or differentially to the monitoring of self-generated timing behavior.

To address this question, we adapted the time production and error-reporting task from our previous study (Kononowicz et al., 2022) and used a within-subject pharmacological inactivation approach to selectively inhibit either the OFC or the ACC while rats performed the task. This design allowed us to assess how each region contributes both to the generation of temporally precise actions and to the evaluation of their accuracy.

## Results

### Rats produce time intervals and monitor their temporal errors

Rats were trained to produce a minimum Target duration (T=2.4s) by pressing a lever twice to demarcate the duration. Producing at least T provided access to a reward (Fig. 1A). If time production (TP) was shorter than T, the lever was retracted and presented again after a random interval. Maximal TP giving access to a reward (Tmax) was calculated individually based on the rat’s precision. All rats produced the required time interval T, as all distributions of TP duration peaked after T (Fig.1B, example of two rats). As in the previous study (Kononowicz et al, 2022), the mean of TP was tightly related to SD (LME: ß=0.217, t(135.4)=6.389, p=2.49*10-9, data not shown), showing a nearly flat association with coefficient of variation (CV) (LME: ß=0.0304, t(23.2)=1.462, p=0.157, data not shown).

**Figure 1.**
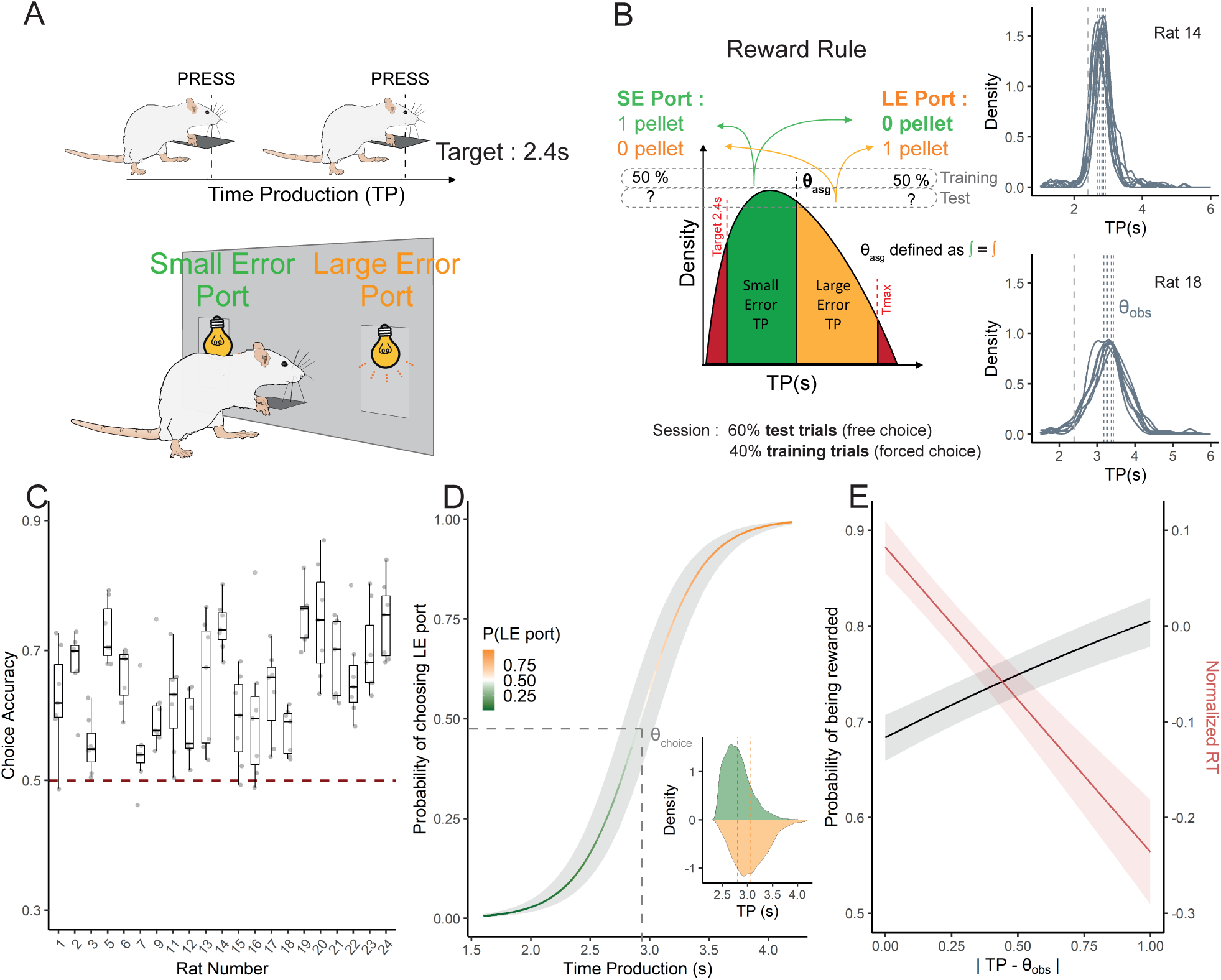
Rats monitor and report their temporal errors. **(A)** Schematic of the operant box design used to train animals, with a lever positioned in the middle of a panel framed by two reward ports. Rats were trained to press twice the lever, with minimum 2.4 s between the two presses to receive a reward, constituting a Time Production (TP). Each port was associated either with « small error » (SE) or « large error » (LE) TP. Reward availability was signaled by the port lit, depicted by the light bulbs. Either one (training trials) or both ports (test trials) were lit immediately after a 2.4 s <TP <Tmax was terminated. Reward delivery (one pellet) was triggered by rats’ nose-poke in the reward port depending on the reward rule depicted in B. **(B)** Inset plots depict TP performance as density plots for rat 14 and rat 18, in test phase (sessions with 60% of test trials) before rats underwent cannula implantation. Based on each rat’s distribution, the main figure of the panel illustrates how rewards were allocated to specific segments of the TP distribution. Green represents SE TPs, while orange represents LE TPs. Red indicates TPs that fell outside the reward range. We calculated the threshold (Θ) between SE and LE TPs as a point dividing the green and orange areas with 1:1 ratio. During the experimental session we used Θ computed based on four previous sessions, namely Θ assigned (Θasg). For the purposes of further analysis, we also computed Θ on the basis of current session, namely observed Θ (Θobs). The arrows point to the probabilistic assignment of TP type (SE or LE) to the left and right ports during training trials. On test trials, the food-port assignments remained unchanged, but both ports were lit, and thus, the retrieval of a reward was determined by the rat’s choice. **(C)** Each column presents a boxplot displaying the median and interquartile range (25th–75th percentiles) and individual sessions (points) choice accuracy for a given rat across all sessions with 60% of test trials. All rats consistently chose above the chance level represented by the horizontal red dashed line, indicating their ability to monitor their temporal errors. (**D**) GLME model fitted to single test trials data. As the curve ascends, TP duration increases together with the probability of choosing LE port, which is in line with the results presented in panel C. Insert plot illustrates the probability density distribution over TPs as a function of rats’ choices. Trials performed by rats as SE are represented in green, while trials performed as LE are depicted in orange. Dashed vertical lines of corresponding colors indicate the mean of each distribution. The SE distribution (orange) is noticeably shifted relative to the LE distribution (green), clearly indicating that rats’ reports were made based on their TPs. We calculated a third threshold, choice Θ (Θchoice) as the TP for which rats have chosen at 50% one port or the other. **(E)** Models fitted to absolute TP distance to the Θobs during single test trials. The distance to Θobs was used as a proxy of confidence. When the TP is farther from Θobs, choices become more certain in line with decision-making models assuming that accrual of evidence is faster from a discrimination threshold. As confidence increases, the likelihood of receiving a reward (black) increases, and response times (RT, red) decrease, indicating that rats make faster and more accurate decisions when they are more confident.

In the crucial part of the task, rats were trained to associate the precision of their TP to one of two options by classifying it as Small Error (SE) productions (Fig.1B, green) and Large Error (LE) time productions (Fig. 1B, orange). We defined SE and LE time productions by calculating a threshold θ for each rat as the TP median of the successful trials in the last 4 sessions (Fig. 1B). Successful trials were defined as trials that fell in the rewarded range, trials in which time production was above the target T and below the outrange criteria Tmax. In order for the rats to learn the rule, we exposed them to training trials in which the port corresponding to their time production category (SE or LE) was lit in 50% of the cases allowing them to link the presence or absence of reward in a given port to their performance in time production. On so-called test trials, both ports were lit up, allowing the rat to choose according to the rule described above, and be rewarded in case of a correct choice. In line with the self-monitoring hypothesis, during these test trials, accuracies in choosing the rewarded ports, which were conditional on a given produced duration, were significantly above the chance level for all of the individual rats (all rats Wilcoxon test: *V*>28, *p*<0.025, *one sided*), as well as as a group (fig. 1C; LME: *ß*=0.152 *t*(20)=10.59 *p*=1.19*10-9). As the test trial accuracy neglects a continuous link between choice and TP precision, we verified that the rats showed an increased probability of choosing LE port as a function of TP (Fig. 1D; GLME *ß*=3.852, *z*=41.20 *p*<10-15). As in the previous study, due to nonstationarity of time production, the history of preceding TPs was justified in the model (AIC=11741) and had an impact on choice behaviour (GLME ß = 2.335, z=10.07 *p*<10-15). However, current trial time productions contributed the most to choice accuracy. Rats were thus continuously assessing their TP to make their choice.

However, we still do not have access to the representation of the rule they are using to categorize their TP. To further characterize rats’ performance, we distinguished three different θ that could be computed by them. The first, termed “assigned θ” (θasg), represents the value assigned by the experimenter to each rat based on the TP distribution from its previous sessions, establishing the reward assignment rule. The second, labeled “observed θ” (θobs), indicates the value determined by applying the reward assignment rule considering the TP distribution in the current session (fig1B). The third, named “choice θ” (θchoice), represents the decision limit at which the rat could not distinguish whether the TP value corresponded to a small or large error trial, that is, the TP value for which it had the same probability to classify the trial as a SE or LE trial (fig 1D). These parameters are based on test trials only. θasg is capturing the actual time value of the threshold that the rat can access through memory of the previous sessions, whereas θobs is corresponding to a theoretical threshold value based on its actual performance during one particular session. Previous work of our team (Nur Bilgin et al., 2026) showed that session choice accuracy is increasing when deviation between θchoice and either θobs or θasg is decreasing, suggesting that rats are using a representation of these parameters to guide their choices at a session level.

To capture trial difficulty in an analogous way to stimulus strength in classical decision-making paradigm, we then expressed single trial TPs relatively to θobs the session-specific threshold computed from that session’s TP distribution. Trials with TPs close to θobs were harder (more ambiguous for categorization), while those farther from θobs were easier. Consistent with this interpretation, the probability of correct choice increased linearly with the trial-to-trial absolute distance between TP and θobs (Fig.1E; GLME: *ß*=0.448 z=6.052 *p*=1.43*10-9), suggesting that rats were basing their choice strategy on a continuous evaluation of their TP. Although rats experience θasg, session-to-session variability in TP means θobs better captures the effective difficulty of each trial.

This hypothesis is further reinforced by the analysis of the normalized Response Time (RT) relative to the same assessment of difficulty. RT were defined as the time the rat was taking to poke in the food port after TP terminated. RTs decreased linearly with the trial-to-trial absolute distance between TP and θobs such that easier trials were executed faster (Fig.1E; LME: *ß*=-0.287 t(19010)=-9.485 *p*<10-15). Altogether it suggests that RT can be used as a proxy of rat’s confidence.

### Different contributions of ACC and OFC to action timing

To investigate the neural bases underlying timing and temporal error monitoring, we infused muscimol, an agonist of GABA-A receptors, to inhibit selectively one of two regions of the prefrontal cortex, ACC or OFC, during the task. Rats underwent cannula implantation and were retrained once consistent error monitoring behaviour was observed. We then alternated infusions of muscimol and control conditions across sessions in a within-subject design, with all subjects completing at least four pairs of sessions. Control sessions comprised saline infusions and sham sessions (identical procedures without infusion), which were grouped together as their TPs did not differ significantly (LMM: *ß*=0.0637, *t*(9.6)=0.827, *p=*0.428, *see Methods)*.

We hypothesized that the ACC could be primarily involved in error monitoring without impacting timing performance. Indeed, we found that, in the ACC group, TPs were not impaired (*ß*=0.00567 t(16960)=0.554 p=0.58) as seen from overlapping density distribution under control and muscimol 2mM (Fig. 2A). The scalar property of timing behaviour is critical for evaluating timing and error monitoring, such that timing variability is preserved across conditions with respect to the timed interval. To assess this property between control and muscimol, we investigated the relationship between the mean and the width of the TP distribution. There was no significant difference in scalar property considering either the standard deviation, or the coefficient of variation between the two conditions (CV: *ß*=-0.011634 *t*(786)=-1.100 *p*=0.27; SD: *ß*=-0.00834, *t*(787)= - 0.268 *p*=0.788). Contrary to the ACC, inhibiting the OFC impaired the rat’s ability to produce time intervals in a dose-dependent manner, with almost no temporal control under the highest 2mM dose (Fig. 2B). Statistical analyses comparing TP under each dose of muscimol to the control condition indicates a disruption of TPs from 0.8mM of muscimol (LMM: 0.5mM *ß*=-0.03289, *t*(7067)=-1.147, *p*=0.251; 0.8mM: *ß*=-0.09629, t(6748)=-3.421, p=0.000627; 1mM: *ß*=-0.2231, *t*(6467)=-8.133, *p*=4.96e-16, 2mM: *ß*=-0.7226 *t*(11390)=-12.575, *p*<2e-16). In addition to shorter TPs, their distribution increased as well, even from the lowest dose of muscimol (LMM: 0.5mM *ß*=0.11534, *t*(653)=2.348 *p*=0.0192; 0.8mM: *ß*=0.43775 *t*(653)=8.912, *p*<2e-16; 1mM: *ß*=0.53219 *t*(652)=11.414 *p*< 2e-16, 2mM *ß*=1.07826 t(653)=10.391 p< 2e-16), also affecting the relationship between TP mean and TP width (LMM: CV 0.5mM *ß*=0.011170 t(649)=0.547, *p*=0.58486; 0.8mM *ß*=0.092187, *t*(617)=4.564 p=6.05e-06; 1mM *ß*=0.054605 t(626)=3.252, *p*=0.00121; 2mM *ß*=0.079864, *t*(662)=2.627 p=0.00882; SD 0.5mM *ß=*0.09927 t(644)=1.438, *p*=0.1510; 0.8mM *ß=*0.53645, *t*(597)=7.882 p=1.53e-14; 1mM *ß=*0.35504 t(608)=6.273, *p*=6.72e-10; 2mM *ß=*0.51632, *t*(663)=5.019 p=6.69e-07).

**Figure 2.**
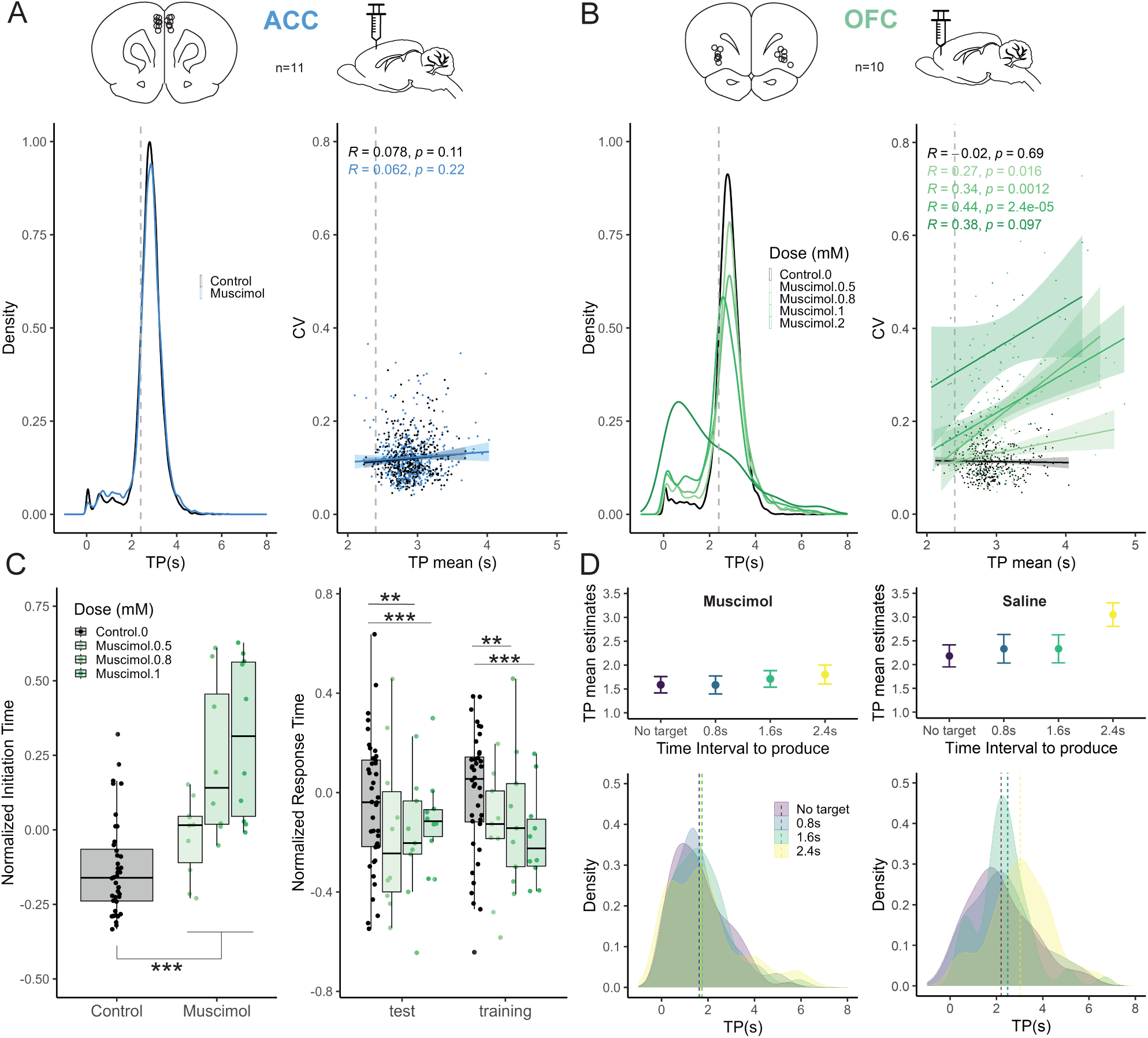
Time Production is impaired under OFC inhibition but not under ACC inhibition. (**A**) Time Production (TP), depicted as probability density over TP, during error monitoring test sessions in control or muscimol condition in the ACC group. TP density was computed on all single trials from all individuals. In both conditions, timing behaviour followed Weber’s law for both groups, showing a nearly flat curve between the coefficient of variation (CV) and the mean TP across sessions and bins (n=10, see *Binning procedure*). (**B**) As in A for the OFC group, illustrating dose-dependent impairment of TP by muscimol. (**C**) Initiation Time (left panel) and Response Time (right panel) under control and increasing doses of muscimol infused in the OFC. The OFC inhibition shortens the length of RT in training and test trials but increases the length of initiation times. (**D**) Timing performance in the control experiment with reduced target duration T (no target, 0.8s, 1.6s, 2.4s). The left and right columns show performance for rats infused in OFC with muscimol 2mM or in control condition for the four Ts tested. TP estimates originate from the model where TPs were fitted with different T. TP increase along with increasing T is only seen in saline condition, demonstrating that rats did not adapt their TP to increasing T under muscimol.

Dose-dependent effects of muscimol were evident in multiple aspects of behavioral performance (Fig. 2C). Normalized initiation time, that is, the interval between lever insertion and the first press initiating a time production, increased progressively with dose, as indicated by LMM estimates (0.5 mM: ß=0.0369, t(5686)=3.745, p=0.00018; 0.8 mM: ß=0.1414, t(5763)=14.542, p < 2e-16; 1 mM: ß=0.1425, t(5457)=14.831, p < 2e-16; 2 mM: ß=0.3500, t(12020)=12.912, p < 2e-16). Normalized RT showed a corresponding progressive decrease with increasing dose, particularly at the higher concentrations (0.5mM *ß*= -0.002154 *t*(7150)=-0.049 *p*=0.96125, 0.8 mM ß=-0.126, t(7404)=-2.844, p=0.0045; 1 mM: ß=-0.254, t(7312)=-5.603, p=2.18e-08). These LMM results indicate a systematic consistent relationship between muscimol dose and behavioral measures, supporting the interpretation of a dose-dependent effect with an opposite effect on initiation time and RT.

When infused with muscimol at 2mM, the rats ceased to perform the task after 20 trials on average. To determine whether this drop reflected motor deficits, motivational changes, or impaired timing production, after the rats had completed all error monitoring sessions, we conducted one additional session in which they were exposed to successive FR1 and FR2 tests under muscimol or saline (Fig. 2D; 2 mM, n = 5 muscimol, n = 2 saline). Rats performed normally in FR1, collecting all rewards, and in the first FR2 block without a minimum interval requirement (no target), they successfully executed the required two lever presses. These results rule out gross motor or motivational deficits.

However, in all the FR2 blocks, muscimol-treated rats consistently produced ∼1.6 s intervals regardless of the minimum interval requirement (no target, 0.8, 1.6, and 2.4 s) indicating a reduced ability to adapt their timing (muscimol: 0.8 s ß=-0.004053, t(425)=-0.022, p=0.983; 1.6 s ß=0.123, t(424)=0.706, p=0.481; 2.4 s ß=0.215, t(360)=1.079, p=0.281). In contrast, saline-treated rats adjusted their intervals according to the threshold, reaching ∼3.0 s for the 2.4 s condition. Importantly, rats were able to complete a final FR1 session even after stopping presses during FR2 blocks.

Disruptions in the OFC have been shown to correlate with increased impulsivity, related to unplanned reactions. According to the impulsivity hypothesis, we expected a dose-dependent decrease in initiation times and response times. However, contrary to the impulsivity hypothesis, we observed an increase in initiation times, which is not aligned with this perspective. As we excluded the possibility of motor impairment we suggest that the effects in OFC signify timing related impairments.

### ACC inhibition impairs self-evaluation of time production

We hypothesized that ACC is primarily involved in temporal error monitoring without impacting timing itself. In line with it, we showed that mean and width of the TP distribution were not significantly different between conditions in the ACC group (see Fig 2A.). To evaluate how inhibition of ACC could alter monitoring of time intervals, we modelled rats choices during test trials as a function of their TP and trial history (GLMM : *Error Judgment ∼ (TP + TP history) * Muscimol*) as we did to characterize choice accuracy before muscimol infusion. To determine the extent of trial history to take into account, we ran a permutation test which demonstrated that the eleven previous TP were contributing to the current one (see supplementary). Based on this test, we decided to use the mean of the last 10 trials as TP history.

The probability of choosing LE Port based on current TP was altered under muscimol condition. Specifically, an overestimation bias (GLMM: intercept effect ß=2.88, z=3.74, p=1.81e-4) indicated that rats judged their TP as longer under muscimol (Fig. 3A) than in the control condition. Interestingly, there was no significant effect on the slope of the logistic curve, suggesting it did not change how rats relied on their evaluation of TP, but rather the evaluation itself (GLMM: slope effect ß=-3.05, z=-1.4, p=0.16). The overestimation bias is also corroborated by the frequency of choosing LE port, which increased under muscimol (Fig. 3B upper part LMM: ß=4.63e-2, t(279)=3.36, p=8.91e-4). Moreover, choice accuracy (Fig.3B, lower part, binned sessions) was diminished in SE port under muscimol (LMM: ß=-0.05165 t(909)=-2.207 p=0.0276), whereas there was no effect of muscimol in the LE port. Choice accuracy during control sessions was similar for both ports (LMM: ß=-0.01462 t(11)=-0.197 p=0.847). These results align well with an overestimation bias, because if rats were more prone to evaluate their TP as longer, they would more often wrongly classify SE TP as LE TP.

**Figure 3.**
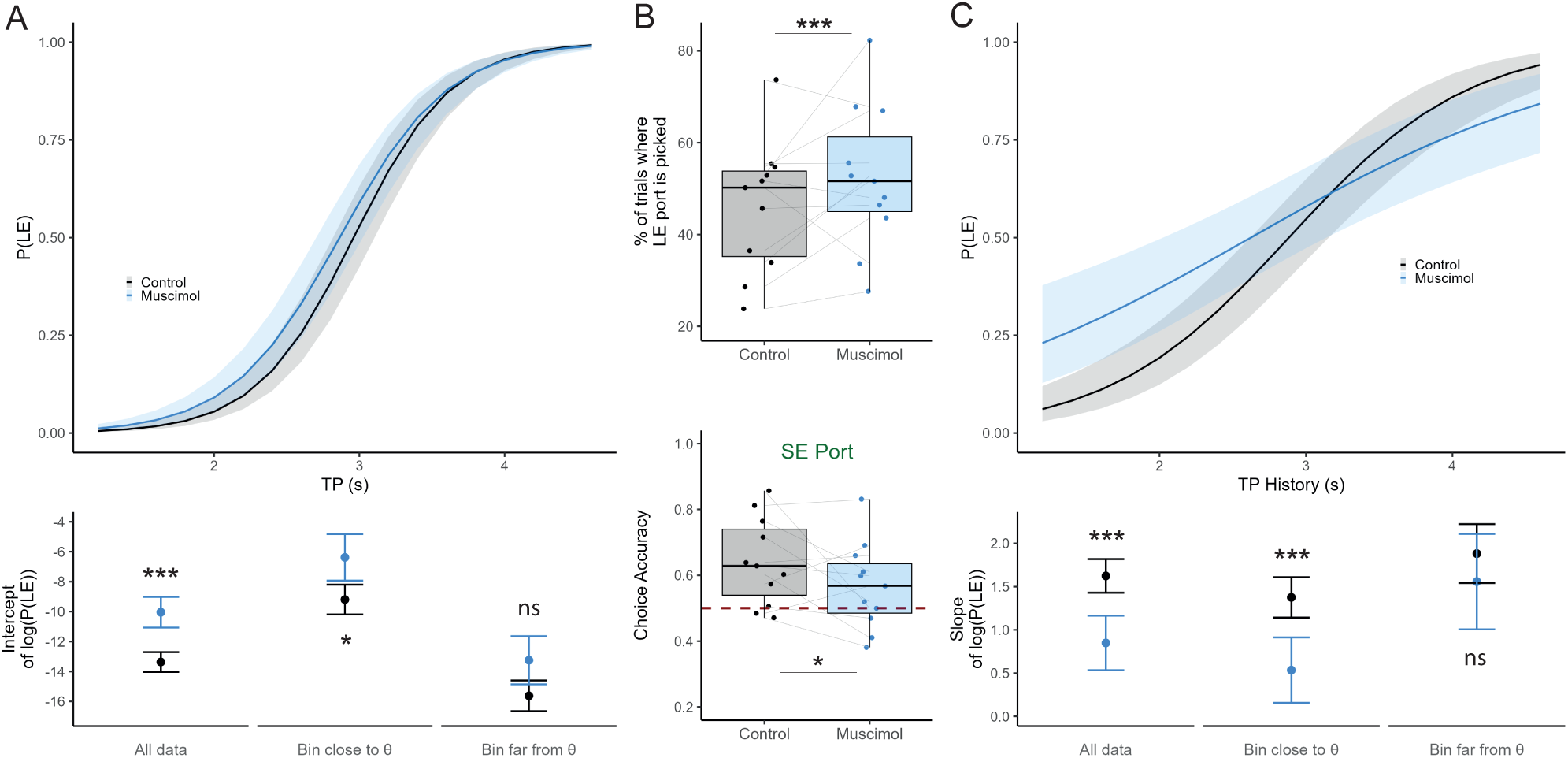
Altered self-evaluation of Time Production under ACC inhibition. (**A**) Model fitted to test trials under control and muscimol conditions featuring the contribution of the undergoing TP to rats’ choices. As the curve ascends, TP duration increases, along with the probability of choosing LE port. However, under muscimol, for the same TP, rats tended to choose LE port more often. Bottom: Beta estimates of intercept for both conditions for all tests trials, trials close and trials far to decision boundaries. (B) Top: Proportion of trials where LE is picked during test trials. Bottom: Choice accuracy for SE trials. (**C**) The same model as in panel A but displaying the effect of trial history. Rats relied on the mean of the last 10 TP to make their choice in a similar, yet less pronounced, fashion. Under muscimol, TP history is contributing less to the probability of choosing LE port. Bottom: Beta estimates of slope for both conditions for all tests trials, trials close and trials far from decision boundaries.

**Figure 4.**
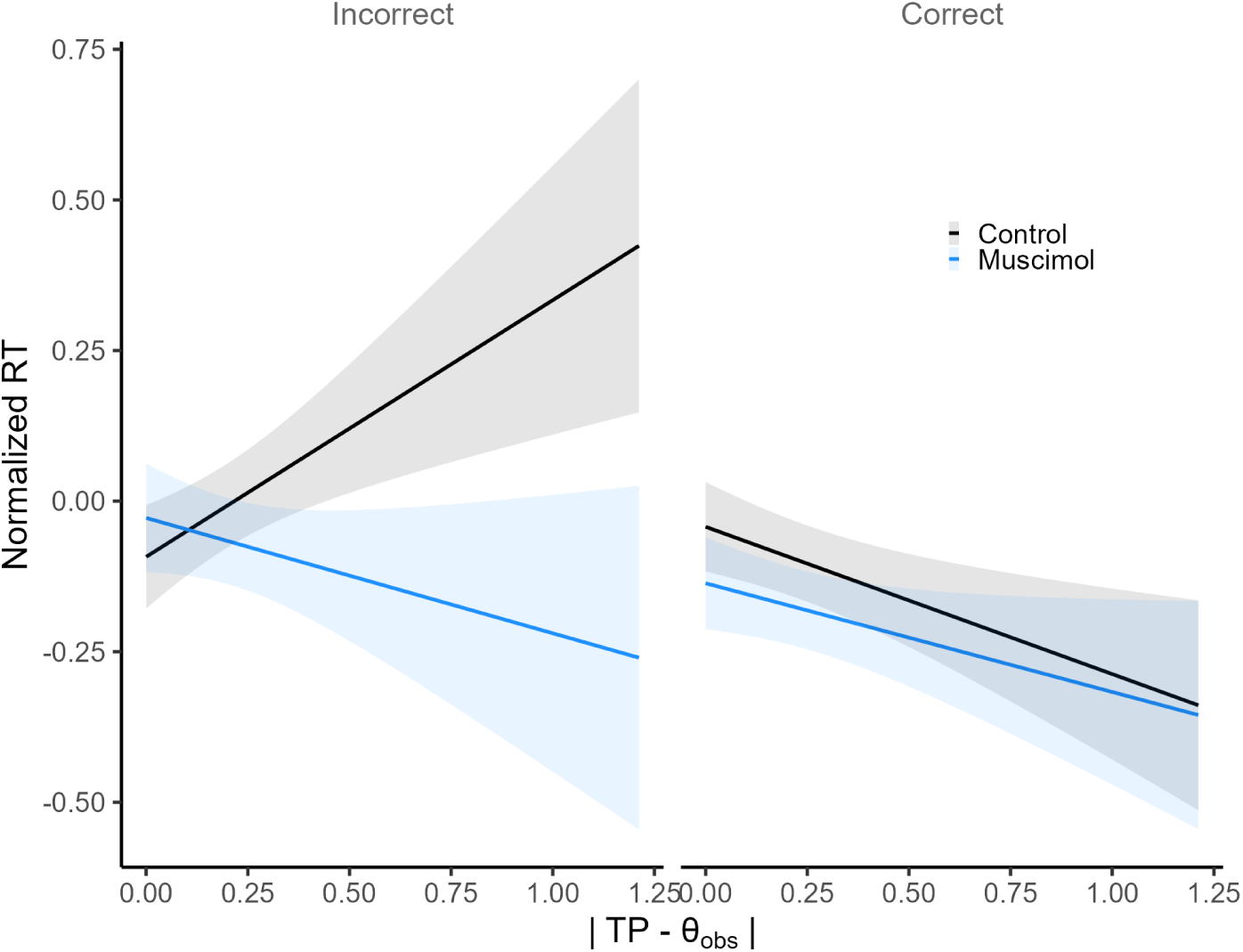
Rats show overconfidence in incorrect easy trials under muscimol. Normalized RT were predicted from a model run on test trials only including single trial distance to, trial correctness, and drug condition. Predicted RTs decreased as confidence increased in correct trials with no effect of muscimol. On the contrary, on incorrect trials, normalized RT increased as confidence increased in the control condition with on opposite effect under muscimol ACC inactivation produced overconfident responses in trials that were actually incorrect.

Yet, this shift in rats’ TP evaluation could originate in a serial correlation of TP toward longer TP due to an impaired trial-to-trial adaptation, especially as ACC has been implicated in behavioural flexibility (Elston et al. 2019). Further analyses show that, even if adaptation was indeed impaired under muscimol, it was not enough to explain by itself overestimation bias (see supplementary).

To analyze whether the impact of muscimol was specific to trials where choices were more difficult to make, we binned the data based on the distance to the θchoice, calculated as the point of subjective equality (PSE) of the logistic curve (see Fig. 1D) for each rat and condition to analyze the effects separately on trials close or far to the rat’s decision boundaries. Muscimol had a significant effect only on the choices made close to θchoice, where having an accurate representation of the task variables is necessary for an optimal choice (Fig. 3A bottom; GLMM: close bin: intercept effect ß=2.42472, z=2.057, p=0.0397; far bin: intercept effect ß=2.2104, z=1.780, p=0.0751). We also observed that ACC inhibition attenuated the impact of trial history on choice accuracy (Fig. 3C; GLMM: slope effect ß=0.627, z=-2.58, p=9.95e-3), suggesting that rats were relying less on the recent history of their past TP to make their choice. This effect was also visible only in trials close to the θchoice (GLMM: close bin: slope effect ß=-0.69991, z=-2.394, p=0.0167; far bin: slope effect ß=-0.2566, z=-0.595, p=0.5519). Interestingly, muscimol did not alter how rats were using their memory of past sessions (see supplementary). These results suggest that inhibiting ACC alters rats’ evaluation of their TP error based on task variables capturing the ongoing session statistics.

### ACC is involved in self confidence in rats decision making

We next examined the impact of ACC inhibition on rats’ confidence in their timing decisions, focusing exclusively on test trials. Trial difficulty was quantified using single-trial distance to Θobs, which served as a proxy for confidence, with larger distances indicating higher confidence. For correct, rewarded trials under control conditions, normalized RT decreased as a function of distance to Θobs reflecting higher confidence and accuracy on “easier” trials. Interestingly, on incorrect, non-rewarded trials, the relationship was reversed: RT increased as distance to threshold increased, suggesting that errors on easy trials may be associated with miscalibrated confidence, consistent with prior observations of the “hard-easy” effect in decision-making (Merkle et al 2009) (LMM: distance β=0.341, t(7725)=2.62 p=0.0088, interaction between distance and incorrect β= -0.539, t(7737) =-3.447 p=0.000569). Critically, ACC inactivation altered this pattern. The interaction between distance to threshold and muscimol treatment was significant (β= –0.453, t(7718)=-2.46, p=0.014), indicating that rats became overall overconfident under ACC inhibition. Furthermore, the three-way interaction with error status (distance × incorrect × muscimol) was also significant (β=0.442, t(7723)=1.97, p=0.049), suggesting that this overconfidence was particularly pronounced on trials where the animal’s timing was incorrect.

These results demonstrate that the ACC contributes to self-monitoring and confidence evaluation in timing decisions. Inactivation leads to a dissociation between actual performance and subjective confidence: rats overestimate the likelihood of a correct choice even on trials that are objectively easy but performed incorrectly. This pattern could arise from different processes of the evaluation process : a reduced drift rate on incorrect trials, leading to less reliable evidence accumulation or an impaired evidence accumulation or an altered decision threshold, producing biased choices towards LE with inflated confidence.

## Discussion

We present the initial evidence supporting the hierarchical and time-independent inference of temporal errors in the ACC. The hierarchy manifests itself in the disruption of error monitoring, which is independent of timekeeping. An alternative model suggests that both processes may be encoded in overlapping neural populations. We propose the existence of a read-out mechanism responsible for inferring temporal errors. This work introduces a novel paradigm and significant original findings for the models of performance monitoring and time perception. When combined with statistical modeling, this data promises to offer valuable insights into ACC function and contributes new insights into neural systems, combining timing with performance monitoring.

### OFC timing, impulsivity and error monitoring

OFC is increasingly recognized for its critical role in decision-making processes at multiple levels, and its role in carrying confidence signals (Lak et al.2014) made it a good candidate for timing error monitoring. Yet, in our task, rats with OFC inhibition were not able to produce the target duration, suggesting a role for OFC in time production upstream to its evaluation.

Interestingly, OFC is known to be involved in evaluating the value of delayed rewards, which relates to timing. For instance, studies have shown that disrupting activity in OFC can impair performances in a delay-discounting decision-making task, leading to a preference for immediate rewards over larger, delayed ones (Mobini et al., 2002, Khani et al., 2014). Moreover, inhibiting OFC with muscimol decreased performances of rats in a waiting time task, and ramping neuronal activity in the OFC was predictive of their waiting duration (Xiao et al 2016). Similar results were obtained in an internally-driven behavioural task where mice had to hold a lever for a minimal amount of time in a self-paced manner (Cazares et al 2022). Transient disruption of lateral OFC after the press initiation prevented the encoding of ongoing press duration which resulted in a drop in performances. OFC could thus have an inhibitory control on cognitive impulsivity through integration of action timing and valuation to guide future optimal behaviour. Muscimol infusion in medial lateral OFC in our task could lead to the suppression of this inhibition of impulsivity, leading to shorter unrewarded TP and reduced RT.

However, we also observe an increase in the latency to initiate a new trial (i.e., the latency to first press the lever after its insertion), which does not comply with this hypothesis. Turner et al (2022) showed that lateral OFC lesion impaired action sequencing and increased delays when initiating and terminating sequences, suggesting that the lateral OFC could be essential to correctly load the motor sequences. Indeed, OFC has been hypothesized to encode information about the current state of the task and update this information based on feedback from the environment (Wilson et al 2014). If this cognitive map is altered under muscimol, it could be difficult for the animal to execute and time correctly the chaining of motor actions constituting the two timed lever presses that rats use to produce duration. Through its role in cognitive mapping and state representation, OFC may also contribute to timing error monitoring on top of its role in time production, a process that could not be assessed here, given the requirement of an initial TP in our task design. Indeed, Brockett et al (2024) showed that inhibiting OFC in an inhibitory control task leads to a deficit in value-based decision making along with a failure to inhibit maladaptive impulsive strategy.

### ACC Temporal Error Evaluation and Confidence

The ACC plays a pivotal role in performance monitoring, serving as a critical hub for error detection and cognitive control. Research indicates that the ACC carries out confidence and error signals across various tasks (Stolyarova et al., 2019; Brockett et al., 2020) and recent work further highlights its role in encoding and updating prediction error signals to guide adaptive behavior (Hassani et al., 2024). Moreover, it has been shown that ACC was essential for a rule switch task where monkeys had to distinguish between different levels of performance failures, such as incorrect choices versus misjudgments about task states (Sarafyazd & Jazayeri, 2019). Disrupting the network showed that ACC was working downstream of the dorsomedial frontal cortex and was specifically involved in inferring covert rule switches, suggesting a hierarchical treatment of the error signals in the ACC.

In line with this framework, inhibiting ACC in our task led to an overestimation bias of rats’ own temporal errors without perturbing their temporal production. This dissociation indicates that ACC can encode internal temporal errors independently from timekeeping mechanisms. We propose that ACC functions as a “read-out” system, extracting temporal production statistics from upstream timing signals, likely originating in the OFC, and computing the error amplitude to guide subsequent actions. Notably, under muscimol, rats’ choice thresholds (θchoice), which represent the boundaries for classifying their time productions as small or large errors, were shifted away from the session-specific thresholds (θobs), indicating that ACC inhibition specifically disrupts the evaluation of ongoing performance rather than the generation of temporal intervals themselves.

Further supporting the role of ACC in hierarchical error monitoring, our analyses revealed that ACC inactivation reduced the influence of trial history on choice behavior, particularly for trials near θchoice, where accurate evaluation is most critical. Additionally, rats under ACC inhibition displayed overconfident responses on incorrect trials, as evidenced by faster response times for “easy” choices they misclassified. Together, these results reinforce the view that ACC acts downstream of timing-related signals from OFC, performing a hierarchical read-out that evaluates temporal errors and confidence, while leaving the primary time production mechanism intact.

### Implications for Hierarchical Models of Temporal Control

These findings align with hierarchical models of temporal error monitoring (Sarafyazd & Jazayeri, 2019), in which timing and evaluation are implemented by distinct but interacting processes. In this framework, a primary “timer” generates temporal intervals, while a downstream “reader” evaluates these intervals to infer error magnitude and guide behavior. Our results support this architecture by demonstrating that OFC inhibition disrupts time production, whereas ACC inhibition selectively impairs error evaluation without affecting timing itself.

Within this organization, temporal information generated upstream is used by ACC to compute internal error signals and confidence estimates. The overestimation bias and altered confidence observed under ACC inhibition reflect a disruption of this evaluative process rather than a deficit in timing. This dissociation highlights a functional separation between the generation and monitoring of internal signals, enabling flexible behavioral adaptation in the absence of external feedback.

### Neural Substrates and Future Directions

While our results establish a functional dissociation between OFC and ACC, the underlying neural dynamics remain to be characterized. We will address this using local field potential (LFP) recordings in both regions during task performance. Human studies show that beta-band activity tracks self-generated time intervals (Kononowicz & van Rijn, 2015), suggesting a candidate neural substrate for temporal representations. Within this framework, OFC may encode temporal information in such dynamics, which are then read out by ACC to compute error and confidence, with inter-regional coordination revealing how these signals are transmitted and evaluated.

## Methods

In the previous study by Kononowicz et al. (2022), the methodologies were described in detail. In this section, we outline the specific procedures and modifications made to the training protocol and primary task. For a visual representation of these protocols, we suggest consulting the supplementary materials available in the earlier publication.

### Subjects

We used 24 male Sprague-Dawley rats (Envigo, France) that were approximately 3 months old at the start of the experiment. Their initial weight was between 240-260g. Rats were housed in groups of 4 and maintained under a 12-hour light/dark cycle, with experiments conducted during the light cycle. Following a week of acclimation in animal housing with unrestricted access to food and water, rats underwent food restriction until their weight reached and was sustained at 85% of their free-fed weight. Daily handling, weighing, and feeding were conducted for all rats throughout the experiment. On days when rats underwent two experimental sessions per day, those sessions were separated by approximately 3 hours, during which they returned to their home cages with access to water. They received food after the second session. The study adhered to the guidelines outlined by the European Communities Council Directive (2010/63/EU) for the care and use of laboratory animals, receiving approval from the French Ministry of Research and the French National Ethical Committee (2013/6).

### Behavioural procedure

Six operant boxes, each with a retractable lever and two food ports (see Fig. 1A) were used for the whole experiment. During 15 sessions of *pretraining*, rats learned to press the lever to activate the light cue in one of the ports where they had to nose poke to get one pellet (45mg dustless grain-based pellets, Bioserv).

Rats were then trained to generate a target duration (T) of 2.4 seconds spaced between two lever presses, where the first press and the second press served as the onset and offset times, respectively. They had to produce a duration of at least T to get access to a reward.

To do so, we first exposed the rats to a T of 0.4s and progressively increased it logarithmically until 2.4s (T: 0.8, 0.6, 1.3, 1.8 seconds). We introduced a new T as soon as 80% of the area under the density function for the TP distribution exceeded the T criterion for two consecutive sessions for one individual (with a minimum of 4 sessions per T). *Duration training* took between 28 and 34 sessions depending on the rats.

Once all subjects achieved the criterion for correct performance on the final target duration (2.4 seconds), a boundary to prevent too long durations was set to 2 * σ(TP) and referred to as Tmax. Rats were not reinforced in trials in which their TP exceeded Tmax. To distinguish these too-long non rewarded trials from too-short non rewarded trials, a tone (4 kHz, 1 second, 65 dB) was played after the TP was terminated. This procedure ensured that animals were precise in their TP and prevented the rats from using strategies that involve stretching their timing distributions to perform better in the subsequent error monitoring phase. This *precision training* lasted 10 sessions.

During *Error Monitoring Training,* a threshold (Θ), which was used for categorization of small error (SE) and large error (LE) TPs was estimated based on individual TP distributions (Fig. 1B). The main alteration from the method used in Kononowicz et al. (2022) was the adoption of a symmetric reward rule. Here, the rats were rewarded with 1 pellet for both Small Error (SE) and Large Error (LE) trials, instead of 2 and 1 pellet, respectively, in the previous study. Thus, in the present study, the point dividing the green and orange areas in the TP distribution was determined with a 1 to 1 ratio to calculate Θ, which will be named Θassign (Fig. 1B). We opted for a symmetric reward rule, as previous research showed contribution of OFC (Padoa-Schioppa, 2011; Schoenbaum et al., 2011) and ACC (Chen et al., 2024) in the valuation process. Therefore, any alterations of choice accuracy could be ascribed to subjective value alterations of two ports. Individual thresholds (Θ) were computed for each animal, with updates triggered when rats deviated from the optimal ratio by a factor of 3 (exceeding the lower boundary of 0.16 or upper boundary of 1.5). Thresholds were recalculated when the updated boundary was exceeded on two consecutive sessions. Figure 1B illustrates the contingencies assigning TP length to a reward port. On a given trial, after TP generation, if TP fell between T and 2*σ(TP), either the left or right port was randomly lit. For trials for which TP fell within the SE criterion (SE trials, green area), animals received 1 pellet in the “SE port” or 0 pellet in the “LE port.” For LE trials (orange area) the reward delivery assignment was opposite (1 pellet in the “LE port”, and 0 pellet in the “SE port”). We considered that animals understood the rule when they began to display lower response times when visiting a rewarded port (Wilcoxon, p<0.5) showing anticipation for the reward. Error Monitoring Training lasted for 28 sessions.

In *Error Monitoring Test* sessions, to assess the rats’ ability to report temporal errors, we used the same trial structure as in the error monitoring training, but we introduced test choice trials in addition to the training trials (Fig. 1B). On test trials both ports were illuminated, allowing the animals to choose based on the previously established rule. In these test trials, rats had the opportunity to receive a reward in either port contingent on their TP. We progressively increased the proportion of test trials from 20% to 60% (by 10% step) to optimize the overall trials count for the analysis of choice behaviour, given the limited number of sessions possible under muscimol infusion. We checked for any alteration of choice accuracy or overall willingness to undergo the task at each proportion tested. Each session allowed the rats to earn up to 100 pellets, with a session time limit of 60 minutes. In total, the rats participated in 22 error monitoring test sessions before they underwent surgery. All figures and analyses on data from the test phase before surgery (Fig. 1) were computed from the last 7 sessions at 60% of test trials.

Rats were then assigned to ACC or OFC groups, such that there was the same number of rats from the same operant box and cage in each group and the median of the choice accuracy during test phase was the same in each group.

### Surgery

All surgeries were performed using aseptic stereotaxic procedures under isoflurane gas anesthesia (2.5-3%). Rats were placed in a stereotaxic frame, the scalp was then incised and retracted and the skull was leveled such that bregma and lambda were in the same horizontal plane. To ensure future stability, three surgical steel screws (PlasticsOne) were implanted into the skull, and then steel guide cannulas (PlasticsOne, 26 gauge) were placed, aiming bilaterally at the OFC (coordinates from bregma AP=+3.8mm, ML=+/-2.6mm, DV=-3.6mm from the skull surface; Paxinos and Watson, 2007) or at the ACC (AP=+1.2mm, ML=+/-0.6mm, DV=-1.2mm). Dental cement (Super-Bond C&B, Sun Medical, Japan) was applied to stabilize the implants. Dummies protruding 0.5 mm from the guide cannulae’s tips were inserted to prevent any clogging. Animals were given seven days to recover from surgery prior food restriction and subsequent behavioural testing. Behavioural measures of discomfort and conditions of the wounds were monitored daily during this period.

### Pharmacological Inactivations

Temporary inactivation was accomplished by localized infusions of the type A **γ**-aminobutyric acid (GABAA) receptor agonist, Muscimol (Sigma Aldrich). On each testing day, the dummies were substituted with injectors protruding 1 mm from the guide cannulae’s tips. After ensuring proper injector placement, Muscimol (2mM, 1mM, 0.8mM or 0.5mm diluted in saline) or sterile saline (NaCl 0.9%, Lavoisier) was injected in OFC or ACC. 0.2 μL (for ACC) and 0.4 μL (for OFC) of infusion fluid was delivered per site at a rate of 0.1 μL/min via a 10μL Hamilton syringe infusion pump (Harvard Apparatus). The injections were controlled by observation of the movement of a small air bubble in the tubing to verify the fluid’s flow. Once the infusions were finished, the injectors were retained in position for 2 minutes to allow diffusion. Behavioural testing started 30 min after the infusion. For the Sham infusions (2 sessions), we replicated the whole procedure at the exception of the activation of the pump. We infused the rats on 8 consecutive days with an alternance of Saline/Sham and Muscimol infusion (4 sessions each). Muscimol was always infused at 2mM for ACC rats and doses were decreased across the session from 2mM to 0.5mM for the OFC rats.

### Histology

Rats were euthanized following the last testing session with an overdose of pentobarbital. Brains were extracted and fixed in 10% formalin acetate for three days, followed by 30% sucrose for 5 days. They were then sliced in 40 μm coronal sections. Cannulae placements were checked by examining slices under a light microscope. Only rats with injector tips within the targeted region were included in statistical analyses.

### Data processing

Custom written scripts in R (version 2022.02.2+485; R Core Team; 2008) were employed for all analyses. Before surgery, individual session datasets were excluded only in cases of malfunctions of lever/pellet distributors or rat’s unwillingness to perform the task. No individual dataset (per rat and per session) was excluded from the 60% test trials sessions. After the surgery, we added two more criteria : wrong location of the injectors tips and perturbed infusion. In total, 16 individual datasets (per rat and per session) were excluded out of 176 datasets after surgery.

We cleaned the data such that we excluded the trials where (TP- µ(TP)) > 3*σ(TP).

To eliminate the left-tail reflecting impulsive responses and motor errors, only TPs longer than 1.5s were analyzed for Weber’s law check and theta indexes.

We normalized Response Time (RT) and Initiation time to be able to compare rat’s performances. Initiation time was z-scored per rat, and RT was z-scored per rat and per port.

### Binning procedure for infusion sessions

In order to maximize the data points for the Weber’s law check (Fig. 2) during the infusion sessions, we binned the data across trial numbers to determine 10 values of mean (µ) and standard deviation (σ) of TP per session and per rat. All the bins were constituted of a minimum of 15 data points.

Choice accuracy was calculated on binned data in all the analyses of the infusion sessions. We binned the test trials per port type (SE or LE) such that we obtained 6 values of choice accuracy per port type and per rat and session. Depending on the analysis we binned choice accuracy across duration of the session or across TP (expressed as absolute distance to θ).

### Statistical analysis

Data were analyzed using linear mixed-effects models (LMMs) in R (*lme4* v1.1-35.1) to account for repeated observations from the same animals across sessions. P-values for fixed effects were obtained using Type III ANOVA with Satterthwaite approximation of degrees of freedom (*lmerTest* v3.1-3). The mixed-effects model approach was integrated with model comparisons to systematically select the best-fitting model. Model selection was performed using the Akaike Information Criterion (AIC).

To account for the hierarchical structure of the data, we included random intercepts for animals and evaluated sessions nested within animals (1 | rat/session). The nested structure allows the model to capture both between-animal variability and within-animal session-to-session variability, which reduces the risk of inflated Type I error. Random slopes were not included because preliminary analyses showed no consistent variation in treatment effects across animals, and inclusion did not improve model fit. Sessions were re-indexed from 1 to 4 separately for each condition (muscimol and control) to avoid confounding treatment effects with session order.

### Permutation test

To determine how far back a significant relationship existed between current TP n and any TP n-back (Fig. 3B), we built a LMM modeling the relationship between the current TP n and the n-back TP (assessing up to TP n-15). We then shuffled the order of a particular n-back 1000 times within individual rats/sessions and compared the shuffled distribution of beta coefficients to the actual value via permutation test, as described in Schreiner et al, 2022.

### Calculation of task variables (θ)

The first, termed “Assigned θ” (θasg), represented the value assigned to the subject based on the TP distribution from their previous sessions, establishing the reward assignment rule. The second, labeled “Observed θ” (θobs), indicated the value determined by applying the reward assignment rule considering the TP distribution in the current session. The third, known as “Choice θ” (θchoice), represented the decision limit at which subjects could not distinguish whether the TP value corresponded to a small or large error trial.

## References

Akam, T., Rodrigues-Vaz, I., Marcelo, I., Zhang, X., Pereira, M., Oliveira, R. F., Dayan, P., & Costa, R. M. (2021). The Anterior Cingulate Cortex Predicts Future States to Mediate Model-Based Action Selection. Neuron, 109(1), 149–163.e7.

Brockett, A. T., Tennyson, S. S., deBettencourt, C. A., Gaye, F., & Roesch, M. R. (2020). Anterior cingulate cortex is necessary for adaptation of action plans. Proc Natl Acad Sci U S A, 117(11), 6196–6204.

Brockett, A. T., Kumar, N., Sharalla, P., & Roesch, M. R. (2024). Optogenetic Inhibition of the Orbitofrontal Cortex Disrupts Inhibitory Control during Stop-Change Performance in Male Rats. eNeuro, 11(5), ENEURO.0015-24.2024.

Cai, X., & Padoa-Schioppa, C. (2012). Neuronal encoding of subjective value in dorsal and ventral anterior cingulate cortex. Journal of Neuroscience, 32(11), 3791–3808.

Cazares, C., Schreiner, D. C., Valencia, M. L., & Gremel, C. M. (2022). Orbitofrontal cortex populations are differentially recruited to support actions. Current Biology, 32(21), 4675–4687.

Chen, W., Liang, J., Wu, Q., & Han, Y. (2024). Anterior cingulate cortex provides the neural substrates for feedback-driven iteration of decision and value representation. Nature communications, 15(1), 6020.

Cohen, M. X., Heller, A. S., & Ranganath, C. (2005). Functional connectivity with anterior cingulate and orbitofrontal cortices during decision-making. Cognitive Brain Research, 23(1), 61–70.

Dehaene, S., Lau, H., & Kouider, S. (2017). What is consciousness, and could machines have it. Science, 358(6362), 486–492.

Hassani, S. A., Tiesinga, P., & Womelsdorf, T. (2024). Noradrenergic alpha-2a receptor stimulation enhances prediction error signaling and updating of attention sets in anterior cingulate cortex and striatum. Nature Communications, 15(1), 9905.

Hayden, B. Y., Heilbronner, S. R., Pearson, J. M., & Platt, M. L. (2011). Surprise signals in anterior cingulate cortex: neuronal encoding of unsigned reward prediction errors driving adjustment in behavior. Journal of Neuroscience, 31(11), 4178–4187.

Khani, A., Kermani, M., Hesam, S., Haghparast, A., Argandoña, E., & Rainer, G. (2014). Activation of cannabinoid system in anterior cingulate cortex and orbitofrontal cortex modulates cost-benefit decision making. Psychopharmacology, 232(12), 2097–2112.

Kononowicz, T. W., & van Rijn, H. (2015). Single trial beta oscillations index time estimation. Neuropsychologia, 75, 381–389.

Kononowicz, T. W., van Wassenhove, V., & Doyère, V. (2022). Rodents monitor their error in self-generated duration on a single trial basis. Proceedings of the National Academy of Sciences, 119(9), e2108850119.

Kononowicz, T. W., Roger, C., & van Wassenhove, V. (2019). Temporal Metacognition as the Decoding of Self-Generated Brain Dynamics. Cereb Cortex, 29(10), 4366–4380.

Kononowicz, T. W., & van Wassenhove, V. (2019). Evaluation of Self-generated Behavior: Untangling Metacognitive Readout and Error Detection. J Cogn Neurosci, 31(11), 1641–1657.

Lak, A., Costa, G. M., Romberg, E., Koulakov, A. A., Mainen, Z. F., & Kepecs, A. (2014). Orbitofrontal cortex is required for optimal waiting based on decision confidence. Neuron, 84(1), 190–201.

Masset, P., Ott, T., Lak, A., Hirokawa, J., & Kepecs, A. (2020). Behavior-and modality-general representation of confidence in orbitofrontal cortex. Cell, 182(1), 112–126.

Mobini, S., Body, S., Ho, M. Y., Bradshaw, C. M., Szabadi, E., Deakin, J. F., & Anderson, I. M. (2002). Effects of lesions of the orbitofrontal cortex on sensitivity to delayed and probabilistic reinforcement. Psychopharmacology, 160(3), 290–298.

Merkle EC. The disutility of the hard-easy effect in choice confidence. Psychon Bull Rev. 2009 Feb;16(1):204–13. doi: 10.3758/PBR.16.1.204. PMID: 19145033.

Niki, H., & Watanabe, M. (1979). Prefrontal and cingulate unit activity during timing behavior in the monkey. Brain Res, 171(2), 213–224.

Padoa-Schioppa, C. (2011). Neurobiology of economic choice: a good-based model. Annual review of neuroscience, 34, 333–359.

Padoa-Schioppa, C., & Cai, X. (2011). The orbitofrontal cortex and the computation of subjective value: consolidated concepts and new perspectives. Annals of the New York Academy of Sciences, 1239(1), 130–137.

Paxinos, G. and Watson, C. (2007) The Rat Brain in Stereotaxic Coordinates. 6th Edition, Academic Press, San Diego.

Sarafyazd, M., & Jazayeri, M. (2019). Hierarchical reasoning by neural circuits in the frontal cortex. Science, 364(6441), eaav8911.

Schoenbaum, G., Takahashi, Y., Liu, T. L., & McDannald, M. A. (2011). Does the orbitofrontal cortex signal value?. Annals of the New York Academy of sciences, 1239(1), 87–99.

Schreiner, D. C., Cazares, C., Renteria, R., & Gremel, C. M. (2022). Information normally considered task-irrelevant drives decision-making and affects premotor circuit recruitment. Nature communications, 13(1), 2134.

Stolyarova, A., Rakhshan, M., Hart, E. E., O’Dell, T. J., Peters, M. A. K., Lau, H., Soltani, A., & Izquierdo, A. (2019). Contributions of anterior cingulate cortex and basolateral amygdala to decision confidence and learning under uncertainty. Nature communications, 10(1), 4704.

Totah, N. K., Kim, Y. B., Homayoun, H., & Moghaddam, B. (2009). Anterior cingulate neurons represent errors and preparatory attention within the same behavioral sequence. J Neurosci, 29(20), 6418–6426.

Turner, K. M., Svegborn, A., Langguth, M., McKenzie, C., & Robbins, T. W. (2022). Opposing Roles of the Dorsolateral and Dorsomedial Striatum in the Acquisition of Skilled Action Sequencing in Rats. The Journal of neuroscience : the official journal of the Society for Neuroscience, 42(10), 2039–2051.

Wallis, J. D., & Kennerley, S. W. (2011). Contrasting reward signals in the orbitofrontal cortex and anterior cingulate cortex. Annals of the new York Academy of Sciences, 1239(1), 33–42.

Wilson, R. C., Takahashi, Y. K., Schoenbaum, G., & Niv, Y. (2014). Orbitofrontal cortex as a cognitive map of task space. Neuron, 81(2), 267–279.

Xiao, X., Deng, H., Wei, L., Huang, Y., & Wang, Z. (2016). Neural activity of orbitofrontal cortex contributes to control of waiting. The European journal of neuroscience, 44(6), 2300–2313.

