## Supplementary material for "Differential Contributions of Anterior Cingulate and Orbito-Frontal Cortex to action timing and its self-monitoring in rats"

### Supplementary Materials

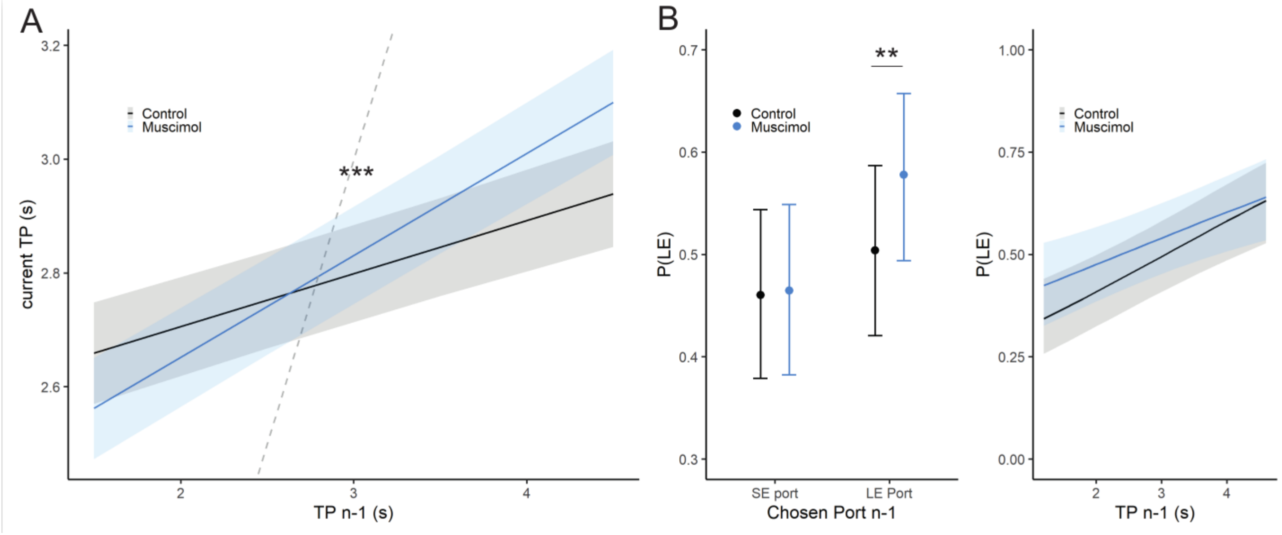


**Supp Fig 1.Inhibiting ACC reduces flexibility** (**A**) Current TP is correlated to the previous TP (TP n-1) showing trial to trial adaptation of TP. Under muscimol, this correlation grows stronger, suggesting a tendency to repeat their TP. (**B**) GLME model fitting the probability to choose LE port to either the last port chosen (left panel), or the previous TP (right panel). In the left panel, we see that under muscimol, rats are more prone to choose LE port if they already picked it the previous trial. In the right panel, probability of choosing LE port increases along the previous time production, a relationship not affected by Muscimol.

Error Monitoring is essential to detect mismatches between expected and actual outcomes, and thus contributes to maintain behavioural flexibility. In rodents, ACC has been implicated in various processes ensuring flexibility, such as decision making, anticipation or adaptation (Elston et al. 2019).

In our task, rats are continuously adapting their TP across a session to match the expected duration in a non-stationary fashion. Yet, rats treated with muscimol in the ACC showed a reduced adaptation from one trial to the next (Supp Fig. 1A). In control sessions, current TP depended positively on the preceding TP (LMM: slope ß=0.093 $t(15730)=8.36$ $p<2\times10^{-16}$) reflecting adaptive trial-to-trial adjustments toward the target duration. Under muscimol, this dependency increased (LMM: interaction ß=0.086, $t(15730)=5.64$, $p=1.77\times10^{-8}$), suggesting that rats were more prone to repeat similar TPs, even if they were too short or too long, thereby adjusting TP less dynamically.

We also observed an alteration in spatial alternation across ports. Under muscimol, the probability of choosing LE port increased following a previous LE choice, whereas it was unchanged following a previous SE choice (Supp Fig.1B, left panel; GLMM interaction between muscimol and previous LE choice: ß=0.28, $z=2.75$, $p=5.88\times10^{-3}$). This pattern indicates reduced alternation, with rats showing an increased tendency to repeat LE choices rather than a global bias toward the LE port. We then evaluated the probability of choosing LE port based on previous TP, and observed that muscimol did not impact the way rats used their assessment of prior TP to guide their choice (Supp Fig1B, right panel, GLMM : muscimol ß=0.281 $z=1.066$ $p=0.286$; interaction muscimol and previous TP  ß=-4.08e-2 z=-0.445 p=0.656) as it would have been expected if the tendency of rats to favor the LE port depended on serial correlations of TPs or a serial pattern of choice repetition.

Thus, inhibiting ACC impacted flexibility through an impaired trial-to-trial adaptation of rat’s TP which cannot explain on its own the overestimation bias of TP we observed during test trials.

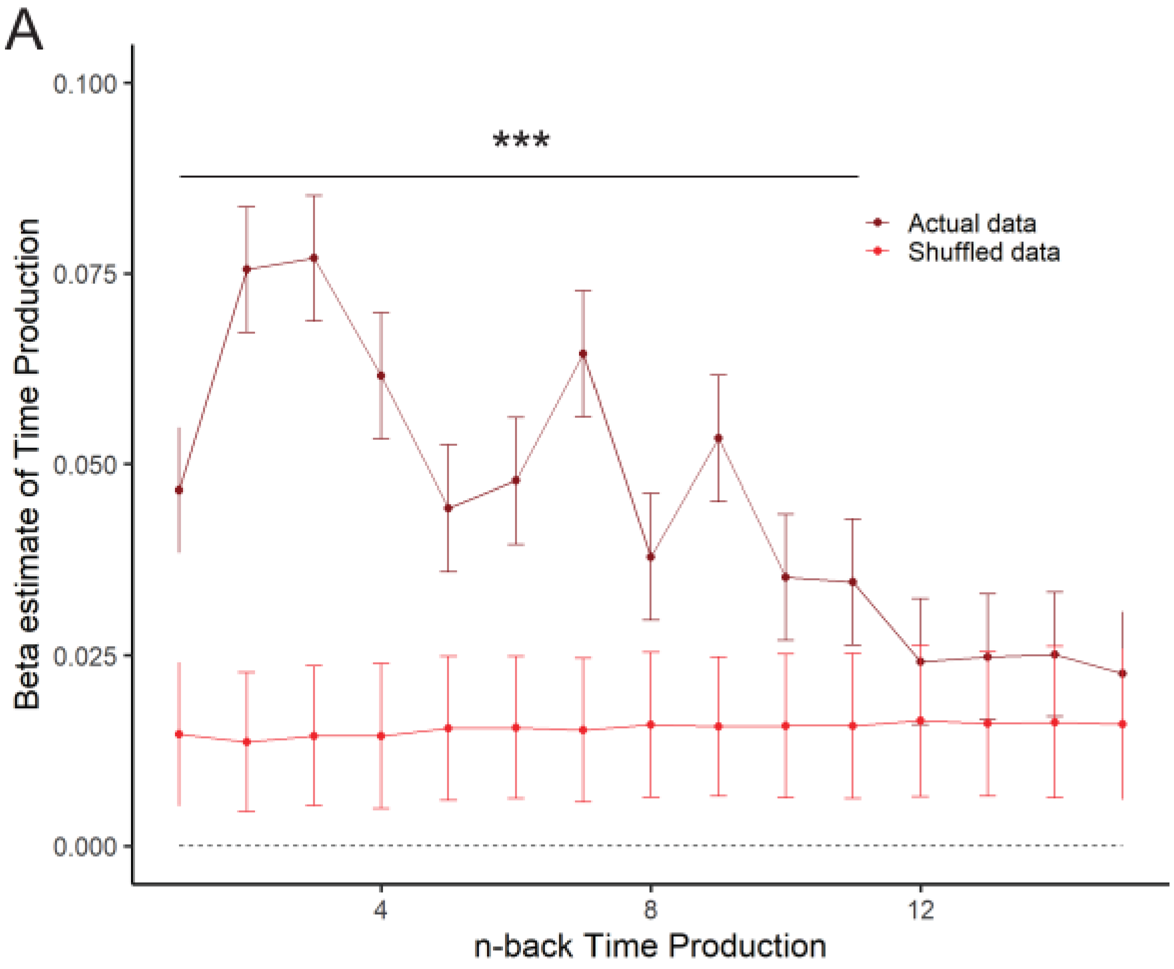


**Supp Fig 2. Past TP are contributing to the current TP.** To evaluate the extent of trial history that rats can maintain in memory, we computed LME model beta coefficients relating current TP to the 15 preceding TP (n - back) for Actual and Shuffled order data. Prior TP beta coefficients up to n-11 differed from the shuffled order data (permutation test $p < 0.001$).

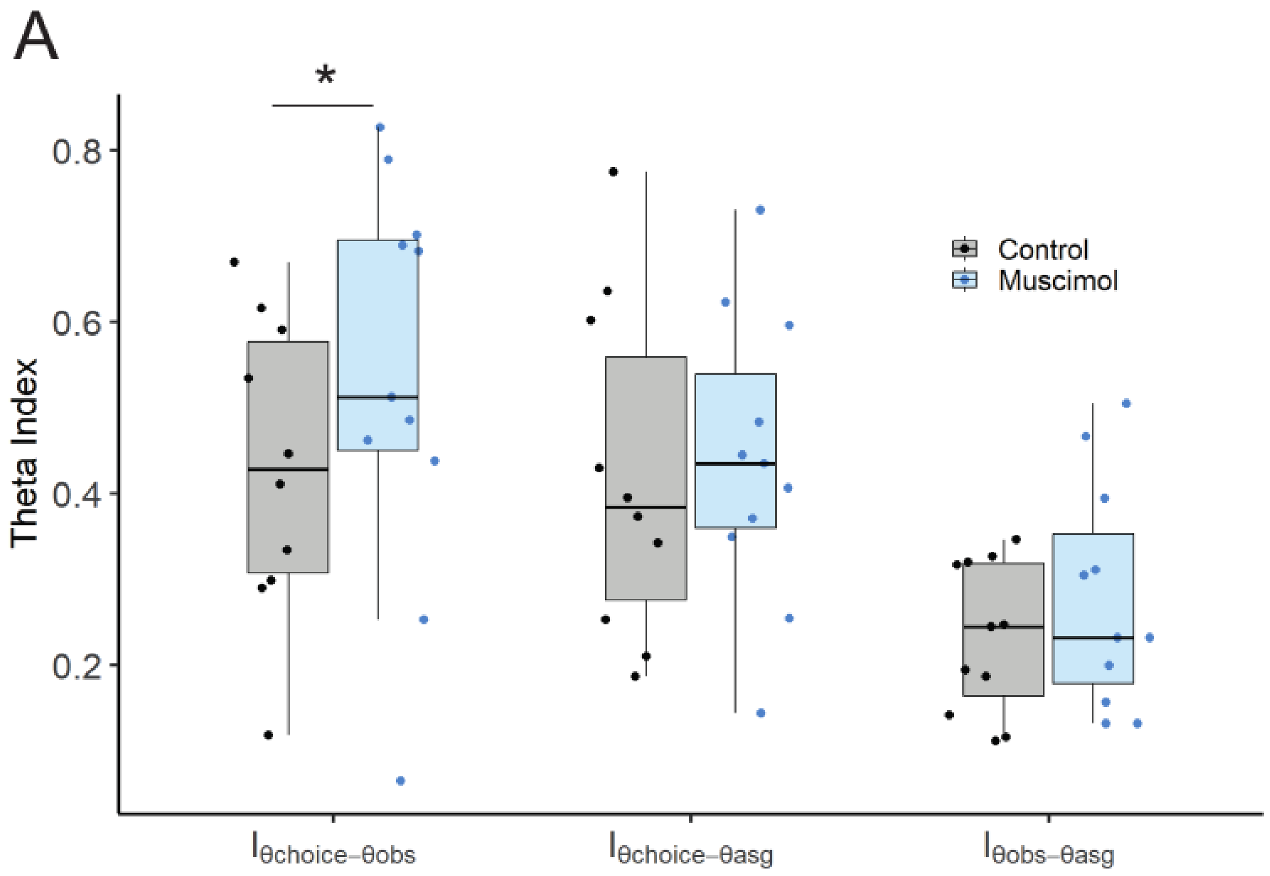


**Supp Fig 3. Inhibiting ACC changes how rats use task variables to guide their choices.** At the session level, task variables were represented by three different thetas : Assigned, Observed and Choice. We computed three indexes expressing absolute distance between thetas. Under muscimol, theta Choice and Observed diverged more under muscimol, suggesting that rats relied less on information from the current session to guide their choices.

Interestingly the distance index between θobs and θasg was smaller than the other two distance indexes, indicating that rats were quite consistent from session to session (LMM: ß=-0.197 t(194)=-3.32 p=0.00109). Under muscimol, the index between θchoice and θobs was slightly increased (LMM: ß= 0.132 t(193)=2.05 p=0.0418), meaning that the decision threshold was further away from the actual threshold than in control sessions. As we showed that timing performance was not impaired under muscimol, it suggests that rats were not using statistics from the ongoing session (current trial TP and memory of previous trials TP evaluation) to the same extent as under control condition to guide their choices. On the contrary, distance between θchoice and θasg was not changed under muscimol, suggesting that rats were still using long-term information from the previous sessions in a similar way.
